# An online dynamic experimental framework to study single neuron responses in humans

**DOI:** 10.1101/2025.07.07.663330

**Authors:** Sunil Mathew, Sofia Dominguez Zesati, Sean Lew, Kunal Gupta, Hernan G. Rey

## Abstract

Single-neuron activity can be recorded from patients being evaluated for neurosurgical treatment of drug-resistant epilepsy who volunteer to have microwires implanted for single cell research that aims to study various aspects of the human brain. These recordings provide an unprecedented window into the human brain to observe how information is encoded and retrieved at the cellular level. Current single cell research paradigms require laborious offline analysis to identify stimuli eliciting neuronal responses and introduce new relevant stimuli in a follow up task. Any delays due to patient availability can compromise the follow up task, risking loss of experimental progress as recording electrodes may move away from previously recorded neurons. This paper introduces a novel experimental framework for online single cell data analysis and dynamic sampling of the stimulus space. It allows for real-time data acquisition and analysis to identify response-eliciting stimuli, introduce new relevant stimuli, and discard those that fail to elicit responses.

## 1 Introduction

Stereoelectroencephalography (SEEG) is a minimally invasive surgical procedure done to identify seizure focus on patients with drug-resistant epilepsy. These patients may choose to participate in single cell research as they present an opportunity to study various aspects of the human brain in both disease and health. Behnke Fried depth electrodes are placed in areas of interest, and they differ from clinical electrodes in that they have microwires protruding from their tip that are able to record single neuron activity. After surgery, the patient is transferred to the Epilepsy Monitoring Unit (EMU), where the electrodes are connected to a neural signal recording system that is able to record the electrical activity of the neurons in the brain. The data is analyzed using various techniques, including spike sorting, which is used to isolate single neuron activity from the recorded signals. Figure 1 shows preoperative MRI images co-registered with post operative CT scans, where electrodes inserted in the amygdala and fusiform areas can be seen.

**Figure 1.**
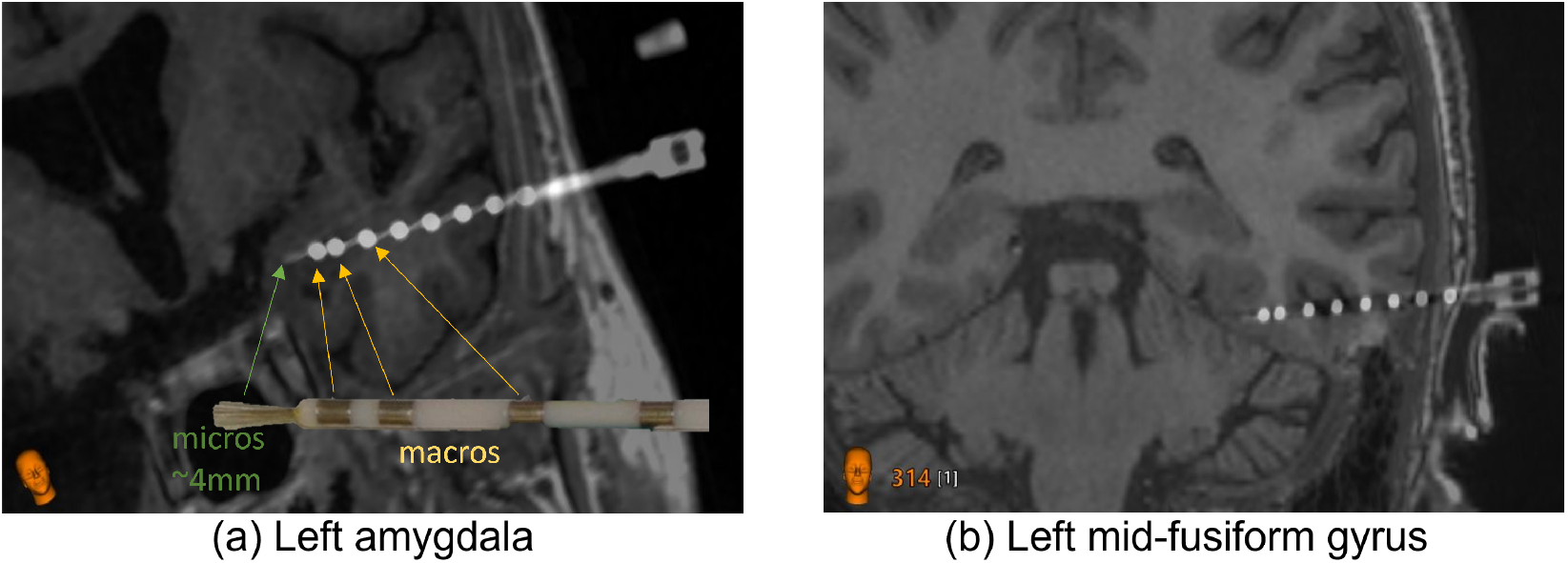
Behnke Fried depth electrodes inserted in the human brain. (a) shows a postop CT coregistered with an MRI. A Behnke Fried depth electrode inserted in the left amygdala can be seen. A real image of the electrode is also placed for reference, showing the macro contacts and the micro wires. (b) Same as in panel (a) but for a trajectory targeting the mid-fusiform gyrus.

Single cell research in humans rely on laborious offline analysis of the recorded signals to identify response eliciting stimuli that are then used to prepare a stimulus set for the follow up experiment. Patient may become unavailable due to a seizure, falling asleep, visitors, lack of interest, or any other unforeseen circumstances, compromising the follow up experiment that was planned. To overcome this issue, in this work we introduce a novel experimental framework for online single cell data analysis in humans that extends the online method introduced by Knieling et al. (2017). While online dynamic paradigms have been employed in animal sensory recordings (Benda et al., 2007; Chambers et al., 2014; DiMattina and Zhang, 2013), our approach is specifically tailored for use in the human invasive recording setting and is applicable to complex, multimodal stimulus domains. The proposed framework not only allows real-time data acquisition and analysis to identify response-eliciting stimuli and discard those that fail to elicit responses, but also introduces new relevant stimuli contingent on the neural responses and predefined rules tailored for the experiment being run. This is made possible due to the advancement of multicore processors and online spike sorting algorithms. The framework is designed to be flexible and can be adapted to conduct experiments that has a block design with any type of stimuli (image, audio, text etc.). The framework is implemented in both MATLAB and Python and is available as open source software.

## 2 System Overview

Figure 2 shows a high-level overview of the system that hosts the online dynamic framework for conducting behavioral experiments. The system comprises both hardware and software components. It has three main processes that form a closed loop: the behavioral task process which delivers the task to the patient (1), the online data acquisition process (3) which gets data from the Neural Signal Processor (NSP) hardware (2), the online process which processes the neural responses and dynamically selects task stimuli (4). The offline local data storage process (5) is run using software provided by the NSP manufacturer (eg: Cerebus, Trellis) for offline data analysis. All processes run on separate threads and communicate with each other via event based messaging during the course of the experiment. Based on the hardware setup specified on the system configuration file, these processes can be hosted on a single computer with no network switches or multiple computers using network switches to communicate between devices as seen in Knieling et al. (2017). This provides flexibility in system design based on available hardware. For example, the data acquisition software provided by the Neural Signal Processor (NSP) manufacturer can be run on a dedicated PC for local storage of the neural signal data for offline processing. A second dedicated PC can be used to deliver the behavioral task to the patient screen (auditory, visual or other) and a third PC maybe used to perform the online neural signal processing that dynamically selects task stimuli based on the neural responses. The system configuration file can be used to select which manufacturer’s NSP is being used (e.g., Blackrock Neurotech, Ripple Neuro), so that the corresponding API calls are made behind the scenes to acquire data for the online processing without needing any additional code change. It will also be used to set the Data Acquisition (DAQ) device being used (e.g., Texas Instruments, Diligent) to provide a digital timestamp (e.g., an image has been sent to the patient screen), or the photodetector being used to detect that an image has been displayed on the patient screen. The experiment is conducted in blocks, with an experiment configuration file specifying how many blocks and how each block is configured (e.g., n_pics=180, n_rep=6, seq_length=30s, isi=0.5s, translates to 18 sequences of 60 pictures each). At the end of every sequence, the behavioral task signals the online process to invoke the neural signal processing pipeline and resumes the task when the online process signals back completion of processing the sequence(1-3secs). At the end of every block, the experimenter is provided with the best responses for review. Selections made by the experimenter then trigger the dynamic selection of task stimuli for the subsequent block. The following sections describe the neural signal processing pipeline and the online dynamic experimental framework in detail.

**Figure 2.**
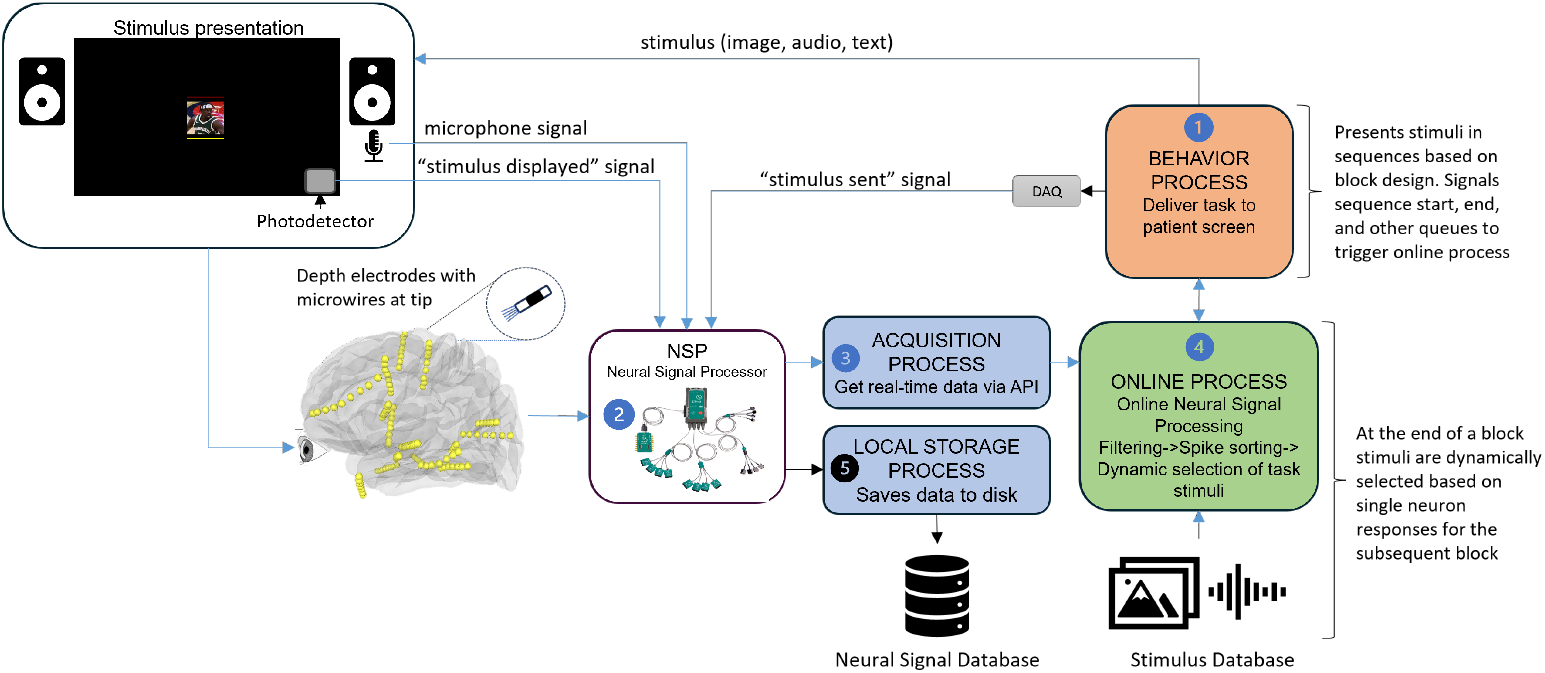
System Overview. (1) The Behavior process presents stimuli to the patient. (2) The neural responses along with signals from DAQ, photodetector, and audio devices are processed by the NSP. (3) The raw data from the NSP is acquired by the online process via the acquisition process. (4) The online process communicates with the behavior process to initiate the online processing pipeline which at the end of a block dynamically selects stimuli for the subsequent block. (5) The local storage process saves the data to disk for offline processing.

## 3 Neural Signal Processing Pipeline

Figure 3 shows the neural signal processing pipeline. It consists of several stages, including power spectrum analysis, bandpass and notch filtering, spike detection, bundle artifact removal, and spike sorting. The power spectral analysis is done just before the start of the experiment to create custom filters for each channel that will be used for the duration of the experiment. The other stages are done online during the experiment by the online process when signaled by the behavioral process at the end of every sequence. The following sections describe each stage of the pipeline in detail.

**Figure 3.**
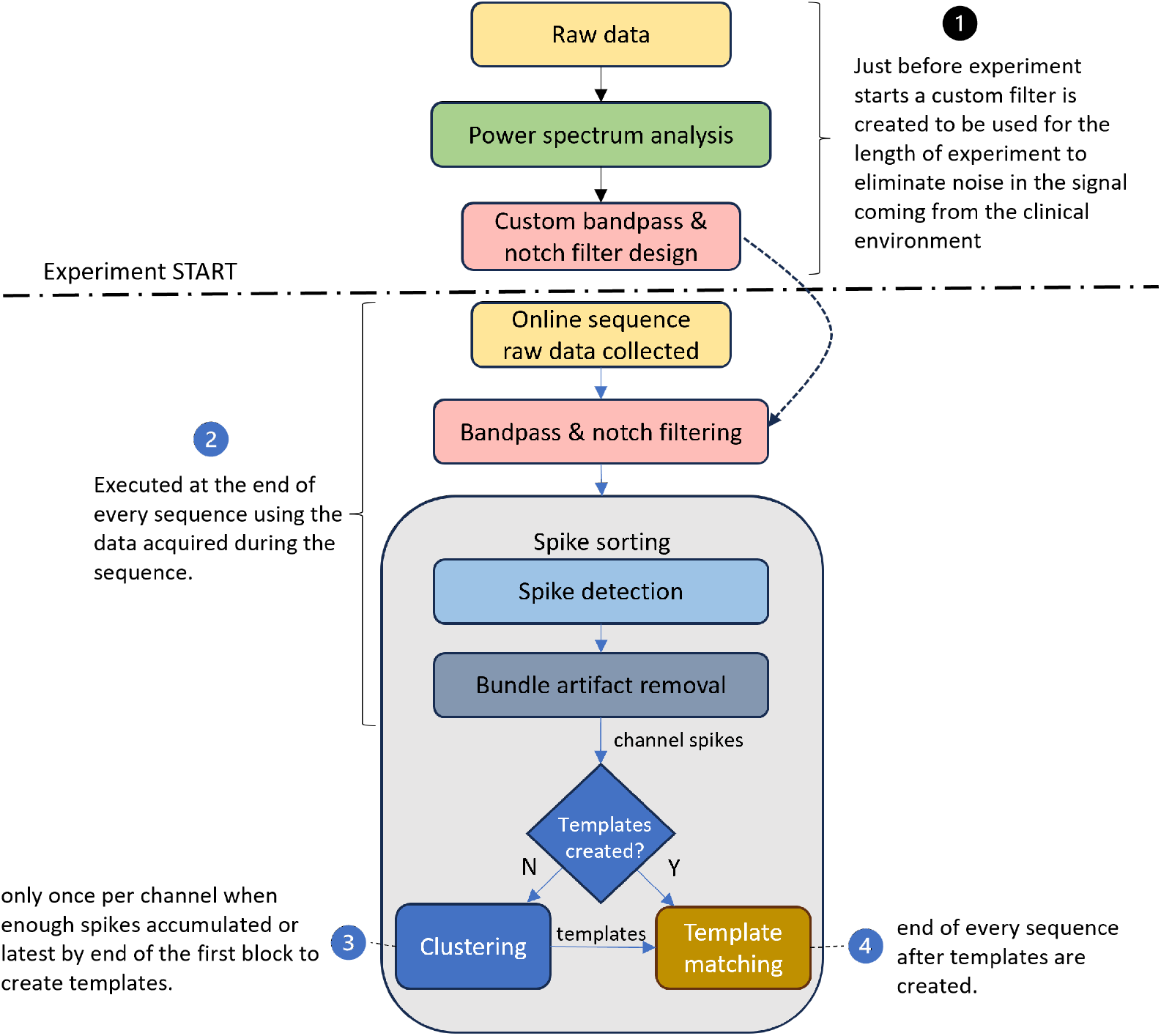
Neural signal processing pipeline. This diagram illustrates the stages of the pipeline, including power spectrum analysis, bandpass and notch filtering, spike detection, bundle artifact removal, and spike sorting.

### 3.1. Power Spectral Analysis, Bandpass & Notch Filtering

Neural signals recorded in a clinical setting are often contaminated by various artifacts, such as line noise (50Hz or 60Hz and its harmonics), mobile phones and interference from bedside monitoring equipment. Power spectrum analysis is an effective tool for identifying and mitigating these sources of noise using filters. The proposed filtering methodology integrates customized notch filtering (for narrowband noise) with bandpass filtering for isolating the signal that pertains to single neuron activity (300 - 3000 Hz). This combination ensures effective noise reduction and signal preservation.

Figure 4 illustrates the power spectrum analysis of a microwire channel that is done just before the start of the experiment to create a custom filter that will be used for the duration of the experiment. The power spectrum reveals the frequency components present in the signal and is calculated using Welch’s method using approximately 2 minutes of data (50 periodograms). To detect spectral peaks requiring attenuation, the computed Power Spectral Density (PSD) is compared to a smoothed version of itself, with an added offset to define the notch detection threshold. Frequencies where the PSD exceeds the notch detection threshold are flagged as candidate notches (black dotted lines in Figure 4 (a)). The amplitude difference between the PSD and the notch threshold is calculated for each candidate frequency. The maximum amplitude difference, A_max_, is determined. The bandwidth for suppressing each significant notch is calculated to be proportional to both the width and relative height of the spectral peak to A_max_. This ensures that wider and stronger peaks will be notched more aggressively with larger bandwidth. Figure 4(c) shows two versions of the filtered signal, the band pass filtered signal (cyan) and the notch filtered version (red) of the bandpass signal. We can see that it is only possible to detect single cell activity in the notch filtered signal, making this an essential step in this processing pipeline i.e. no spikes could have been detected in the experiment without notch filtering. Figure 4(d) shows the spike sorting results obtained with the spikes detected in the notch filtered signal, confirming the good quality of the acquired signal. The following sections describe the spike detection and sorting process in detail.

**Figure 4.**
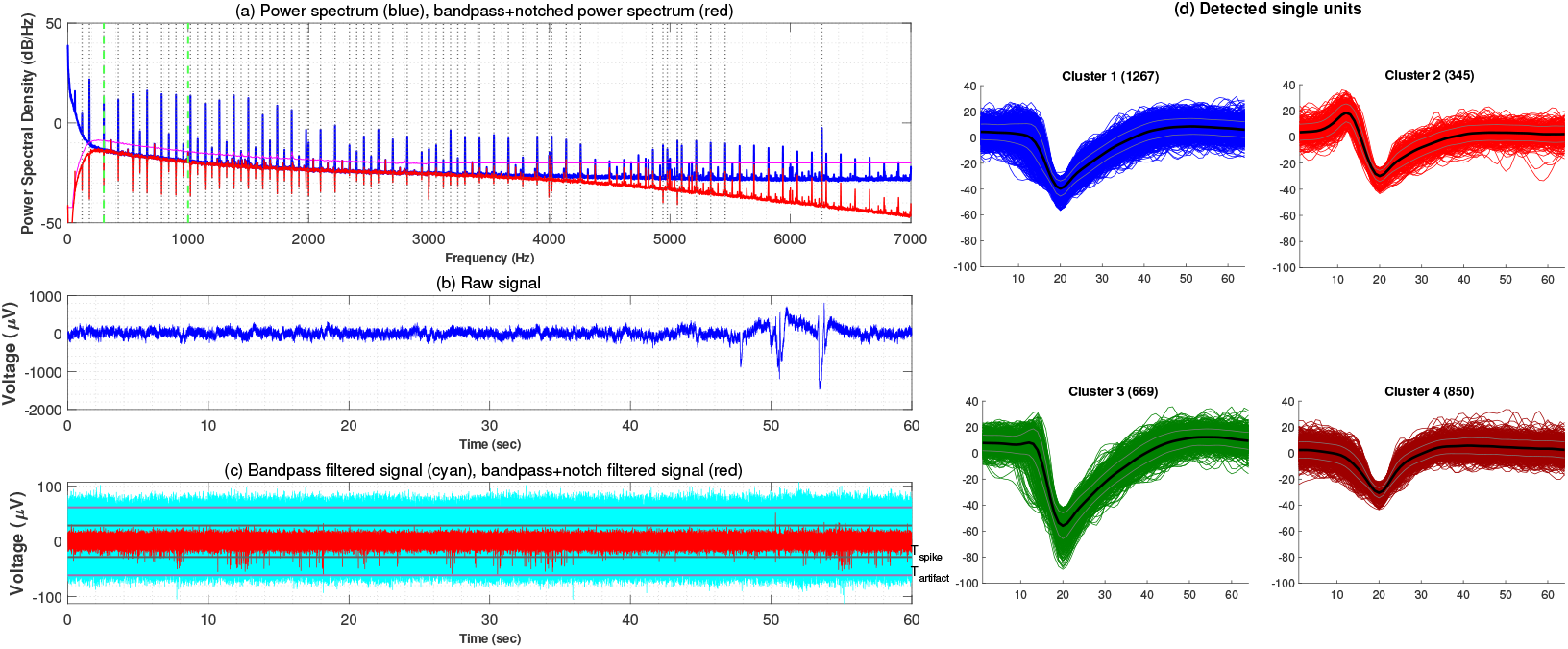
Noise filtering and spike detection. (a) power spectrum of the raw signal (blue) seen in (b), the black dotted lines show the candidate notches that were detected based on the purple notch threshold line and the notch filtered spectrum (red). (c) bandpass signal in cyan (300-3000Hz) and the notch filter applied to it in red. It is evident that the single neuron activity can only be detected after notch filtering the bandpass signal. (d) spike sorting results obtained with the spikes detected in the custom filtered signal.

### 3.2. Spike sorting

Spike sorting is the process of identifying and classifying spikes from putative neurons in the recorded signal Gibson et al. (2012); Rey et al. (2015). The spike sorting process is done in the following steps:

### 3.3. Spike detection

At the end of every sequence, the online process initiates the spike detection process.The detection of spikes and artifacts in neural data relies on the computation of dynamic thresholds, designed to adapt to the noise level of the signal. The noise level in the signal, *σ*, is estimated using the Median Absolute Deviation (MAD), a robust statistical measure of variability,

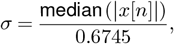

where x[n] represents the input data. The constant 0.6745 scales the MAD to approximate the standard deviation under Gaussian noise assumptions Rey et al. (2015) Two thresholds are defined based on the estimated noise level. The first is the **Spike Detection Threshold (***T*_**spike**_), which is used to identify putative single cell spikes and is defined as,

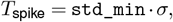

where `std_min` is a predefined scaling factor (e.g., `std_min` = 5). Second is the **Artifact Detection Threshold (***T*_**artifact**_), which is used to detect extreme amplitude signal artifacts from noise, patient movement or other equipment artifacts, this threshold is expressed as,

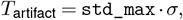

where `std_max` is a predefined scaling factor (e.g., `std_max` = 50).

### 3.4. Bundle artifact removal

Some artifacts that are of below *T*_artifact_ may occur because of patient movement and other external environment factors and can be seen simultaneously on most channels in a microwire bundle. These are identified as bundle artifacts and removed using a bundle artifact removal algorithm. The algorithm works by identifying putative spikes that were detected on most of the channels in a microwire bundle (e.g., 5/8 channels) in a short sliding time window (0.5ms) and removing them from the spike train.

### 3.5. Clustering and Template Matching

As shown in Figure 3, spike sorting is done by the online process using a combination of clustering and template matching for each channel using multithreading. The online process initiates the clustering process when enough spikes have been detected in a channel or at the end of the penultimate sequence of the first block of the experiment, whichever comes first. After clustering is done, templates are created which will be used to classify putative spikes detected in a channel at the end of every subsequent sequence. The clustering and template matching process is done in the following steps:

#### 3.5.1. Feature Extraction

The first step in the spike sorting process is to extract features from the detected spikes. Features are used to create the space where the clustering is going to be performed. The goal is to reduce the dimensionality of the data while preserving the information necessary to enable separability between clusters. Several algorithms can be used to extract features from the spike data, such as principal component analysis (PCA) and wavelet transform. The number of features can be obtained with different methods. For example, a data driven approach introduced in Chaure et al. (2018) is used to identify the wavelet coefficients that will be used. However, hardware specifications for online processing might pose further constraints in the dimensionality of the space where the clustering will be performed.

#### 3.5.2. Online Clustering

A clustering algorithm is used to group the spikes into different clusters based on their features. It is preferable to use an unsupervised clustering algorithm that can automatically detect the number of clusters and finish processing quick enough that it does not add a delay and prolong experiment time. Several clustering algorithms can be used for online spike sorting, such as mountainsort Chung et al. (2017), osort Rutishauser et al. (2006), or combinato Niediek et al. (2016). Again, processor speed and memory for online processing can influence the choice of the clustering method.

#### 3.5.3. Template Matching

After the clustering is done, templates are created for each cluster. Templates are used to classify putative spikes detected in a channel at the end of every subsequent sequence. The templates are created by computing the mean and variability measures for each cluster.. The putative spike is classified as belonging to the cluster with the closest template within a distance limit, otherwise it remains unassigned to any cluster.

## 4 Online dynamic selection of task stimuli

The goal of the online dynamic experimental framework is to identify response-eliciting stimuli, introduce new relevant stimuli, and discard those that fail to elicit responses. This allows for exploration of the stimulus space to understand the neural code better. The experiment is conducted in subscreening blocks as defined in the experiment configuration file. Each subscreening is divided into sequences with a short break (self-paced by the patient) between sequences, and a larger break (<60 sec) at the end of the block while responses are ranked across units and stimuli based on a set response criterion. The best responses are then reviewed, and stimuli can be selected for further exploration of the stimulus space. The framework is designed to be flexible, allowing its use with various types of stimuli (e.g., images, audio, text) and across different experimental paradigms—such as fLoc, a category functional localizer (Stigliani et al., 2015), or tasks involving auditory stimuli (Chambers et al., 2014).

Figure 5 illustrates the dynamic screening process with an RSVP experiment. The first subscreening block is conducted using a subset of the personalized stimulus set that covers the space of all concepts discovered from the questionnaire that the patient fills out prior to the SEEG evaluation. The following sections describe the screening process and dynamic selection of task stimuli in detail.

**Figure 5.**
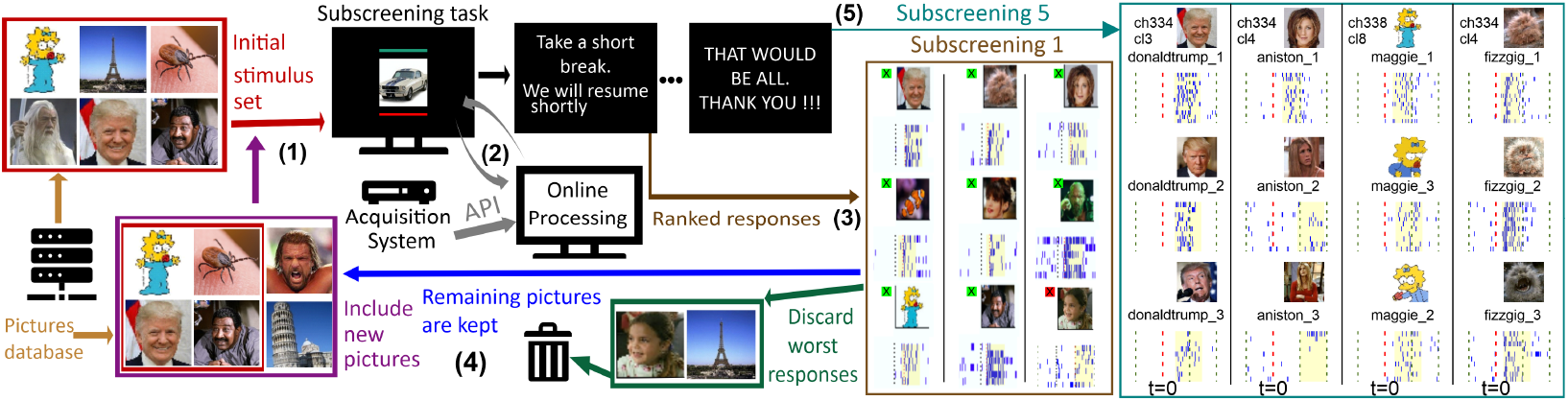
Online and dynamic screening of stimuli. Using an initial set of pictures (180), the first subscreening takes place until all pictures have been shown 6 times each (1). Meanwhile, data is acquired and processed online and in parallel, including spike sorting, without affecting the delivery of the task (2). Then, the patient takes a short break (<60 sec) while responses are ranked across units and stimuli based on a set response criterion. Strong response-eliciting stimuli are then selected for further exploration (3). The worst ranked stimuli are discarded (e.g., bottom 80), allowing for new stimuli to be included (4). This process is repeated for each subscreening. In the sessions presented in this paper, we included two additional pictures of each selected identity. After the last subscreening ends, the rasters for the selected stimuli (and related ones) are quickly obtained (5).

### 4.1. Stimulus Database

The stimulus database is a curated collection of images used in the RSVP experiment. To tailor the content, the patient completes a questionnaire about their interests, hobbies, and personal memories. This is done to increase the chance of identifying response eliciting stimuli during the task (Rey et al., 2015). The patient’s responses are processed to extract key concepts and related items—for example, if a favorite sports team is the Milwaukee Bucks, related items might include Giannis Antetokounmpo and Bobby Portis. Based on these, a personalized set of images are added. In addition to these tailored stimuli, the database also includes widely recognizable images of famous people, popular TV shows, logos, and other culturally familiar references that are generally known, regardless of individual preference.

### 4.2. Rapid Serial Visual Presentation (RSVP) Experiment

RSVP experiments are conducted in blocks, where each block consists of a predefined number of images being presented a predefined number of repetitions to the patient as defined in the block configuration of experiment configuration. Based on the number of repetitions each block consists of *N*_seq_ sequences, given by,

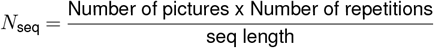

where the sequence length is the number of pictures that can be shown in a sequence. The sequences are created using a block design, where all images that are to be shown in the block are randomized and assigned to sequences to form one repetition, and the procedure is then repeated Nrep times (Rey et al., 2015). The images are shown for a short duration (500 ms) with no gap between them. The task also incorporates an activity to keep the patient engaged, such as a button press when a visual queue like a dot changes color (Allen et al., 2022; Stigliani et al., 2015).

### 4.3. Neural response visualization & review

A visual stimulus like an image needs to be shown several times to the patient to conclude that a neuron fires consistently but only a few times (3-6) might be enough to decide that it is not eliciting a response. Rey et al. (2018) shows that we need to show an image at least 12 times for stable statistical validation and stimulus onset (latency) calculation.

After a stimulus is shown a configured number of times in a block, the responses are ranked, and best responses are visualized as a raster plot along with response metrics and Instantaneous Firing Rate (IFR) as seen in Figure 6. On the left of the stimulus image is its ID, number of trials and the channel number along with the single unit ID. On the right is the z-score and the recording location. The raster plot shows the spike times for all trials over a 1.5 second window (-500ms before stimulus onset to 1000ms after), with each row representing a single trial. The IFR is calculated by computing the histogram of the spikes with short time window (1ms). A threshold based on the baseline IFR (black horizontal line) is calculated, computing mean and standard deviation during a baseline period (e.g., when a blank screen is displayed), and is used to identify the onset (green dotted line) and duration (cyan background) of the response to the stimulus. The spikes during the baseline period and post stimulus period are used to calculate the z-score for estimating the response strength. These response metrics are used to rank the single neuron responses across units and stimuli based on a set response ranking criterion detailed in Rey et al. (2020). Based on the response ranking, the stimuli with the highest response ranking are selected for further exploration, while the stimuli with the lowest response ranking are discarded. Additionally, a stimulus can be marked to be discarded from subsequent blocks as shown in Figure 5.

**Figure 6.**
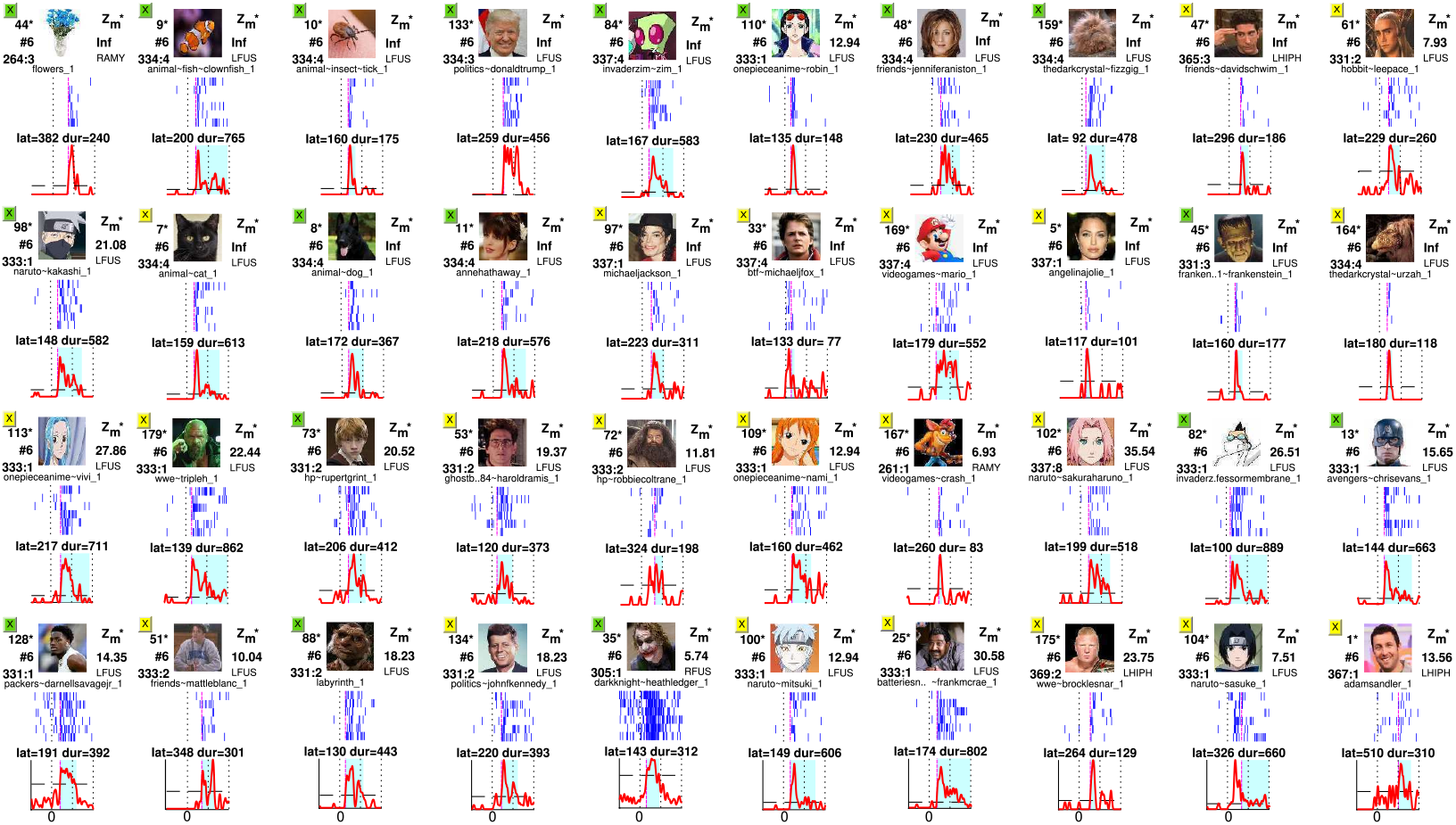
Neural response visualization and review. Responses ranked based on response metrics at the end of the first block of an experiment, shown to the experimenter for review and selection. Selection would be used to dynamically select the task stimuli for the next block of the experiment. The response metrics include the instantaneous firing rate (IFR), onset latency, duration, and z-score. The vertical black dotted lines mark 0, 500ms and 1000ms in the IFR plots and 0ms in the raster plot. The horizontal black dotted line in the IFR plot is the IFR threshold. The magenta dotted line mark the stimulus onset in both the raster plot and IFR. The responses are ranked based on a set response ranking criterion (Rey et al., 2020).

### 4.4. Dynamic selection of task stimuli

The selected stimuli are further explored using a set of predefined task-specific rules to select images from the stimulus database to be used in the subsequent block of the experiment. The rules are designed to promote addition of stimuli that would elicit a response and also to explore the stimulus space based on the task hypothesis. Figure 7 shows the responses from a neuron in the left fusiform area after the second block of the fLoc task (Stigliani et al., 2015). These fLoc stimuli consist of grayscale images with noise-controlled backgrounds and normalized contrast and luminance for several categories, including adult and child faces, bodies, cars, corridors, instruments, limbs, houses, letters, numbers. The exemplar neuron initially responded to three items in the “word” category which were selected after the first block. Associated stimuli were added by the task-specific rules, and all but the last three elicited a response from the same neuron suggesting that this process provides a systematic method to sample the stimulus space. Also, for the first three stimuli, the IFR for the first block (green) and the overall after the second block (red) being consistent suggests that the neuron continues to respond to the stimulus after several repetitions without any signs of stimulus adaptation, which may occur in other species or brain areas.

**Figure 7.**
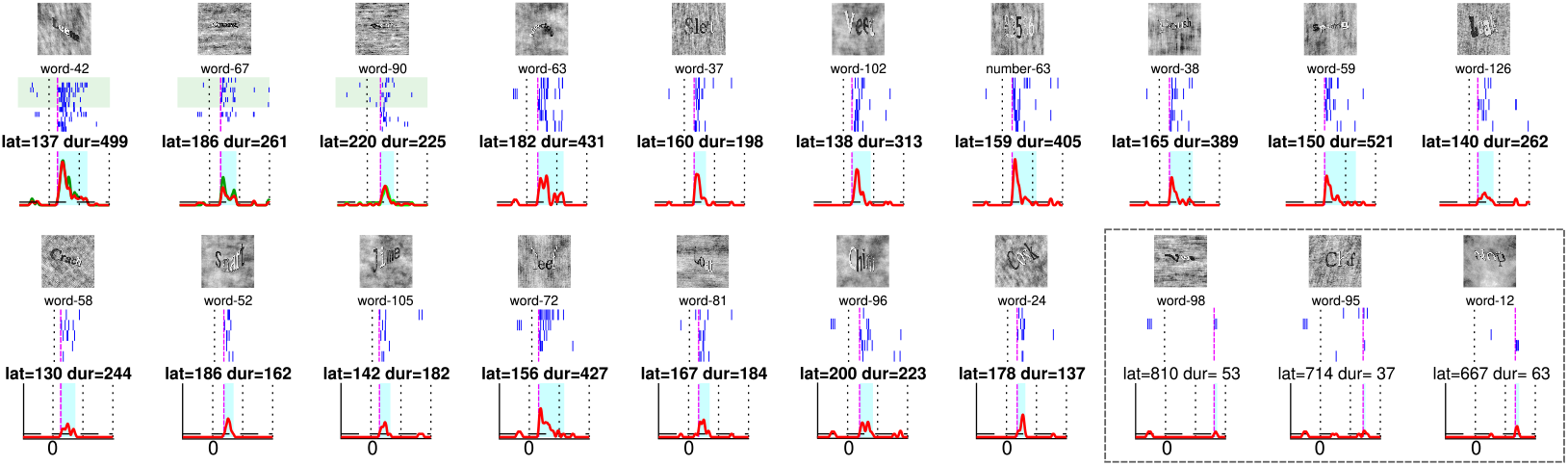
Category localization task stimuli selection. This figure shows responses after the second block of category localization task from a neuron in the left fusiform area. The first three stimuli were selected after the first block, the responses are highlighted in green are the first 6 trials of the first block, the next five rows are responses from the second block. All other stimuli were selected based on predefined rules for the category localization task to be presented in the subsequent block. Except for the last three stimuli, all other stimuli that were added elicited a response from the neuron.

Highly ranked stimuli that were selected and the stimuli that were added based on the rules are protected from being discarded in the subsequent blocks of the experiment until it has been shown for a configurable number of times(*N*_*rep*_ *>*= 12).

## 5 Conclusion

Leveraging the power of modern multicore processors and online spike sorting algorithms, it is now possible to conduct online single cell data analysis in humans to identify response-eliciting stimuli, discard those that fail to elicit responses, and introduce new relevant stimuli based on predefined rules. The framework is designed to be flexible and can be adapted to conduct experiments with any type of stimuli (image, audio, text etc.). The framework is implemented in both MATLAB and Python and is available as open source software.

## Bibliography

1. Allen, E. J., St-Yves, G., Wu, Y., Breedlove, J. L., Prince, J. S., Dowdle, L. T., Nau, M., Caron, B., Pestilli, F., Charest, I., et al. A massive 7t fmri dataset to bridge cognitive neuroscience and artificial intelligence. Nature neuroscience, 25(1):116–126, 2022.

2. Benda, J., Gollisch, T., Machens, C. K., and Herz, A. V. From response to stimulus: adaptive sampling in sensory physiology. Current Opinion in Neurobiology, 17(4):430–436, 2007. doi: 10.1016/j.conb.2007.07.009. Sensory systems.

3. Chambers, A. R., Hancock, K. E., Sen, K., and Polley, D. B. Online stimulus optimization rapidly reveals multidimensional selectivity in auditory cortical neurons. Journal of Neuroscience, 34(27):8963–8975, 2014.

4. Chaure, F. J., Rey, H. G., and Quian Quiroga, R. A novel and fully automatic spike-sorting implementation with variable number of features. Journal of neurophysiology, 120(4):1859–1871, 2018.

5. Chung, J. E., Magland, J. F., Barnett, A. H., Tolosa, V. M., Tooker, A. C., Lee, K. Y., Shah, K. G., Felix, S. H., Frank, L. M., and Greengard, L. F. A fully automated approach to spike sorting. Neuron, 95(6):1381–1394, 2017.

6. DiMattina, C. and Zhang, K. Adaptive stimulus optimization for sensory systems neuroscience. Frontiers in Neural Circuits, Volume 7 -2013, 2013. doi: 10.3389/fncir.2013.00101.

7. Gibson, S., Judy, J., and Markovic, D. Spike sorting: The first step in decoding the brain: The first step in decoding the brain. IEEE Signal Processing Magazine - IEEE SIGNAL PROCESS MAG, 29:124–143, 01 2012. doi: 10.1109/MSP.2011.941880.

8. Knieling, S., Niediek, J., Kutter, E., Bostroem, J., Elger, C., and Mormann, F. An online adaptive screening procedure for selective neuronal responses. Journal of Neuroscience Methods, 291:36–42, 2017. doi: 10.1016/j.jneumeth.2017.08.002.

9. Niediek, J., Boström, J., Elger, C. E., and Mormann, F. Reliable analysis of single-unit recordings from the human brain under noisy conditions: tracking neurons over hours. PloS one, 11(12):e0166598, 2016.

10. Rey, H. G., Pedreira, C., and Quian Quiroga, R. Past, present and future of spike sorting techniques. Brain Research Bulletin, 119:106–117, 2015. doi: 10.1016/j.brainresbull.2015.04.007. Advances in electrophysiological data analysis.

11. Rey, H. G., Ison, M. J., Pedreira, C., Valentin, A., Alarcon, G., Selway, R., Richardson, M. P., and Quian Quiroga, R. Single-cell recordings in the human medial temporal lobe. Journal of Anatomy, 227(4):394–408, 2015. doi: 10.1111/joa.12228.

12. Rey, H. G., De Falco, E., Ison, M. J., Valentin, A., Alarcon, G., Selway, R., Richardson, M. P., and Quian Quiroga, R. Encoding of long-term associations through neural unitization in the human medial temporal lobe. Nature communications, 9(1):4372, 2018.

13. Rey, H. G., Gori, B., Chaure, F. J., Collavini, S., Blenkmann, A. O., Seoane, P., Seoane, E., Kochen, S., and Quian Quiroga, R. Single neuron coding of identity in the human hippocampal formation. Current Biology, 30(6):1152–1159.e3, Mar 2020. doi: 10.1016/j.cub.2020.01.035.

14. Rutishauser, U., Schuman, E. M., and Mamelak, A. N. Online detection and sorting of extracellularly recorded action potentials in human medial temporal lobe recordings, in vivo. Journal of neuroscience methods, 154(1-2):204–224, 2006.

15. Stigliani, A., Weiner, K. S., and Grill-Spector, K. Temporal processing capacity in high-level visual cortex is domain specific. Journal of Neuroscience, 35(36):12412–12424, 2015.

